# Short duration splice promoting compound enables a tunable mouse model of spinal muscular atrophy

**DOI:** 10.1101/2020.08.17.253526

**Authors:** A Rietz, KJ Hodgetts, H Lusic, KM Quist, EY Osman, CL Lorson, EJ Androphy

**Affiliations:** Department of Dermatology, Indiana University School of Medicine, Indianapolis, IN, 46202, USA; Laboratory for Drug Discovery in Neurodegeneration, Brigham & Women’s Hospital, Harvard Medical School, Cambridge, MA, 02139, USA; Department of Veterinary Pathobiology, Bond Life Sciences Center, College of Veterinary Medicine, University of Missouri, Columbia, MO, 65201, USA

## Abstract

Spinal muscular atrophy (SMA) is a motor neuron disease and the leading cause of infant mortality. SMA results from insufficient survival motor neuron protein (SMN) levels due to alternative splicing. Antisense oligonucleotides, gene therapy and splicing modifiers recently received FDA approval. However, early intervention is required for optimal outcomes, and even continuous treatment maybe insufficient to restore full motor function. Although severe SMA transgenic mouse models have been beneficial for testing therapeutic efficacy, models mimicking milder cases that manifest post-infancy have proven challenging to develop. We have established a titratable model of mild and moderate SMA using the splicing compound NVS-SM2. Administration for 30 days prevented development of the SMA phenotype in severe SMA mice, which typically show rapid weakness and succumb by postnatal day 11. Furthermore, administration at day eight resulted in phenotypic recovery. Remarkably, acute dosing limited to the first three days of life significantly enhanced survival in two severe SMA mice models, easing the burden on neonates and demonstrating the compound as suitable for evaluation of follow-on therapies without potential drug-drug interactions.

## Introduction

Spinal muscular atrophy (SMA) afflicts approximately 1 in 6,000-10,000 live births, and half succumb within two years (Verhaart *et al*, 2017). SMA results from insufficient survival motor neuron (SMN) protein. The *SMN1* gene, located on human chromosome 5q13.2, is duplicated and inverted, resulting in the nearly identical *SMN2* gene possessing a nucleotide transition (C→T) in exon 7, causing exon skipping and loss of the terminal 17 amino acids of the SMN protein (Lefebvre *et al*, 1995; Lorson *et al*, 1999; Monani *et al*, 1999). These alternatively spliced *SMN2* transcripts yield a highly unstable protein, SMNΔ7 (Lorson & Androphy, 2000). Only 10-15% of *SMN2* mRNAs produce full-length functional SMN.

SPINRAZA™ (nusinersen), an antisense oligonucleotide, and ZOLGENSMA^®^ (onasemnogene abeparvovec-xioi), an AAV-9 based gene therapy, have recently been FDA-approved for SMA; SPINRAZA™ for all forms of SMA, and ZOLGENSMA^®^ for children under two years. *SMN2* splicing modifiers Risdiplam™ and Branaplam™ are in Phase 3 clinical trials for SMA type I (NCT02913482) and II (NCT02913482) and Phase 2 for type I (NCT02268552), respectively. In SMA type I, clinical trial data indicate reduced lethality and achievement of important motor milestones following intervention with the drugs. Motor functions stabilize in SMA type II patients instead of slowly declining. Risdiplam™ improved the Gross Motor Function Measure scale in SMA type II/III children aged two years and over compared to placebo control (Dangouloff & Servais, 2019). Nonetheless, some patients did not respond to treatment, and there is a strong inverse correlation between the age at which treatment began and efficacy (Dangouloff & Servais, 2019). This highlights the need for co-therapy investigation, as one SMN-modifying agent not be sufficient to completely improve motor skills and disease severity.

The SMN Δ7 SMA (Jackson Lab; Stock Number 005025 FVB.Cg-Tg(SMN2*delta7) 4299AhmbTg(SMN2) 89Ahmb *Smn1tm1Msd*/J) mouse model is most commonly used for testing SMA therapeutics. These mice lack murine *Smn* and express an intact human *SMN2* gene plus SMN2 Δ7 cDNA (Le *et al*, 2005). SMNΔ7 mice develop a severe SMA phenotype with impaired motor function and low body weight with an average life span of 12-13 days (Le *et al*., 2005). A drawback of SMNΔ7 mice is their unfavorable breeding scheme with only 25% of a litter having the SMA genotype. The less-used, slightly more severe ‘Li or ‘Taiwanese’’ SMA mouse model (Jackson Labs; FVB.Cg-Smn1tm1HungTg(SMN2)2Hung/J.) also lacks murine *Smn* and expresses the human *SMN2* transgene(Hsieh-Li *et al*, 2000). These mice display low body weight, gastrointestinal dysfunction, and succumb by postnatal day (PND) 11 (Hsieh-Li *et al*., 2000; Sintusek *et al*, 2016). Their breeding scheme results in 50% of the litter developing the SMA-like phenotype. Following disease progression, both mouse models exhibit necrosis of the ears, tail, and digits due to vascular thrombosis. Similarly, digital necrosis has been reported in infants with severe SMA (Araujo *et al*, 2009; Rudnik-Schoneborn *et al*, 2010). Both mouse models have marked reduction in spleen size (Khairallah *et al*, 2017), which is recapitulated in the less severe Smn^2B-^ mouse model (Khairallah *et al*., 2017) that expresses a knock-in mutation disrupting splicing of endogenous Smn and survives approximately one month (Bowerman *et al*, 2012; Hammond *et al*, 2010; Quinlan *et al*, 2019; Sleigh *et al*, 2011). The C+/+ mouse model (Jackson Lab; FVB.129(B6)-*Smn1tm5(Smn1/SMN2)Mrph*/J) is the mildest genetic model of SMA and exhibits low body weight with very mild impairment of a subset of motor functions, but a normal life span (Osborne *et al*, 2012). This model has been used to investigate i*n vivo* activity of small molecules. These transgenic SMA models are well-characterized and are the go-to standard for therapeutic testing. With the current advances of SMA treatment options, there is now a need to study co-therapies as well as models that more resemble SMA types II and III. To address this need, research has focused on non-genetic approaches with motor dysfunction beginning later in life.

Non-genetic mild SMA mouse models are typically generated in SMNΔ7 or Smn^2B^ mice, though a small number of studies use the 5058 model. Daily administration of *SMN2* splicing modifier SMN-C3 at a suboptimal dose in SMNΔ7 mice induces a milder SMA phenotype (Feng *et al*, 2016) with low body weight and a median life span of 28 days; however, the required daily intraperitoneal injections and oral gavage is a significant burden to the neonatal mice. Other non-genetically induced mild SMA models include suboptimal dosing with AAV9-SMN (Meyer *et al*, 2015), oligonucleotides targeting SMN splicing (Osman *et al*, 2016; Zhou *et al*, 2015), and AAV-9s targeting disease-modifying proteins such as plastin-3 (Kaifer *et al*, 2017) and follistatin (Feng *et al*., 2016). Each intervention presents unique challenges for the studying of co-therapies. Stress resulting from repeated injection in neonatal mice may blunt synergies, and the CMV enhancer/chicken-β-actin promoter used to drive SMN in AAV-9-based interventions may not be consistently activated. Strong and constitutively activated promoters are prone to inactivation due to extensive methylation (Domenger & Grimm, 2019). SPINRAZA™ has an estimated terminal half-life of 135-177 days in the cerebrospinal fluid (CSF) and 63-87 days in the plasma (Neil & Bisaccia, 2019), increasing likelihood of drug-drug interactions. These challenges highlight the need for novel approaches to study co-therapies and to distinguish potential drug-drug interactions.

Our goal was to modify the severe Li SMA mice to a milder SMA mouse model with minimal intervention on the treated newborn that will allow efficacy testing of combinatorial therapies with limited drug-drug interactions. For these studies we used the previously published human *SMN2* splicing modifier NVS-SM2, which promotes exon 7 inclusion and restores normal SMN protein expression, although less efficient in promoting exon 7 inclusion than Branaplam™ (NVS-SM1) at 3 mg/kg in C/+ SMA mice (Palacino *et al*, 2015). The influence of NVS-SM2 on life span in SMA mice has not been reported. Pharmacokinetic analysis demonstrated that NVS-SM2 is readily available in the brain after intravenous (IV) and oral (PO) administration in mouse and rat with Tmax of 3 hours after PO with 3 mg/kg in mice, and treatment induced a 1.5-fold increase in SMN protein the mouse brain (Palacino *et al*., 2015). The advantage of a pharmaceutically induced mild SMA model in the Li mice is their favorable breeding scheme with 50% of their progeny exhibiting symptomatology due to pathologically low levels of the SMN protein.

## Results and Discussion

We synthesized NVS-SM2 and confirmed activity in our previously reported *SMN2* reporter cell assay (Supplemental Figure 1A),(Cherry *et al*, 2012; Cherry *et al*, 2013). NVS-SM2 caused a dose-dependent increase in SMN-luciferase expression up to by 1500%, followed by a decline at ∼3 µM due to cytotoxicity, as indicated by a decrease in renilla-luciferase (Supplemental Figure 1A). To investigate *in vivo* activity, severe SMA mice were generated according to the breeding scheme that produces severe SMA mice (*Tg(SMN2)2Hung*^*tg/0*^; *Smn1*^*tm1Hung/ tm1Hung*^) and heterozygous (Het, *Tg(SMN2)2Hung*^*tg/0*^; *Smn1*^*tm1Hung/wt*^) control siblings as described by Gogliotti *et al*. (Gogliotti *et al*, 2010). Heterozygous (Het) mice express both mouse and human SMN protein, while severe SMA mice express only low levels of human SMN generated from the human *SMN2* transgene. Het and severe SMA neonatal mice were injected subcutaneously (s.c.) with 1 mg/kg NVS-SM2 or vehicle (PEG:PBS, 50:50) once daily for five consecutive days, beginning at PND 2. On PND 7, mice were sacrificed, and tissues harvested. SMN levels across treatment groups were quantified using the human SMN specific monoclonal SMN antibody 2F1. Antibody specificity for human SMN was confirmed in whole brain lysates of Het, severe SMA, and non-transgenic FVB/NJ PND 7 old mice. For comparison, SMN protein levels were also investigated using MANSMA 6, which detects human and mouse SMN protein. The 2F1 antibody did not detect SMN in non-transgenic FVB/NJ mice, whereas MANSMA 6 detected SMN in all three strains (Supplemental Figure 1B). NVS-SM2 treatment increased human SMN protein levels by 4.5-fold in brain (p=0.0005), and 2.5-fold spinal cord (p=0.0355) and muscle tissues in severe SMA mice (Figure 1A-C, Supplemental Figure 2A, B). Human SMN protein levels in muscle of vehicle-treated severe SMA mice were not detectable (Figure 1C). In het control cohorts, we detected higher levels of human SMN in all tissues compared to severe SMA (Supplemental Figure 2B). We hypothesize that the mouse and human SMN proteins expressed in the het control mice are stabilized due to the oligomerization properties of SMN (Lorson *et al*, 1998). NVS-SM2-treatment also increased SMN protein in brain, spinal cord and muscle in Het siblings. This extends the work by Palacino *et al*., who reported a 1.5-fold increase in SMN protein in C/+ mice brain tissues after a single 30 mg/kg NVS-SM2 administration (Palacino *et al*., 2015). These authors found that NVS-SM1 and NVS-SM2 have similar pharmacokinetic properties in mice and rats. However, NVS-SM1 resulted in a greater increase in *SMN2* splicing in C/+ mice than NVS-SM2, while both increased SMN protein in brain tissue at 30 mg/kg oral treatment by 1.5-fold in C/+ mice. In comparison, we show that a 30-times lower dose of NVS-SM2 administered daily subcutaneously on five consecutive days increased brain SMN protein by 4.5-fold severe SMA and Het mice. These differences in SMN expression may be due to the administration method or the mouse models used. Additionally, NVS-SM2 induction of *SMN2* splicing may occur more slowly than with NVS-SM1.

**Figure 1.**
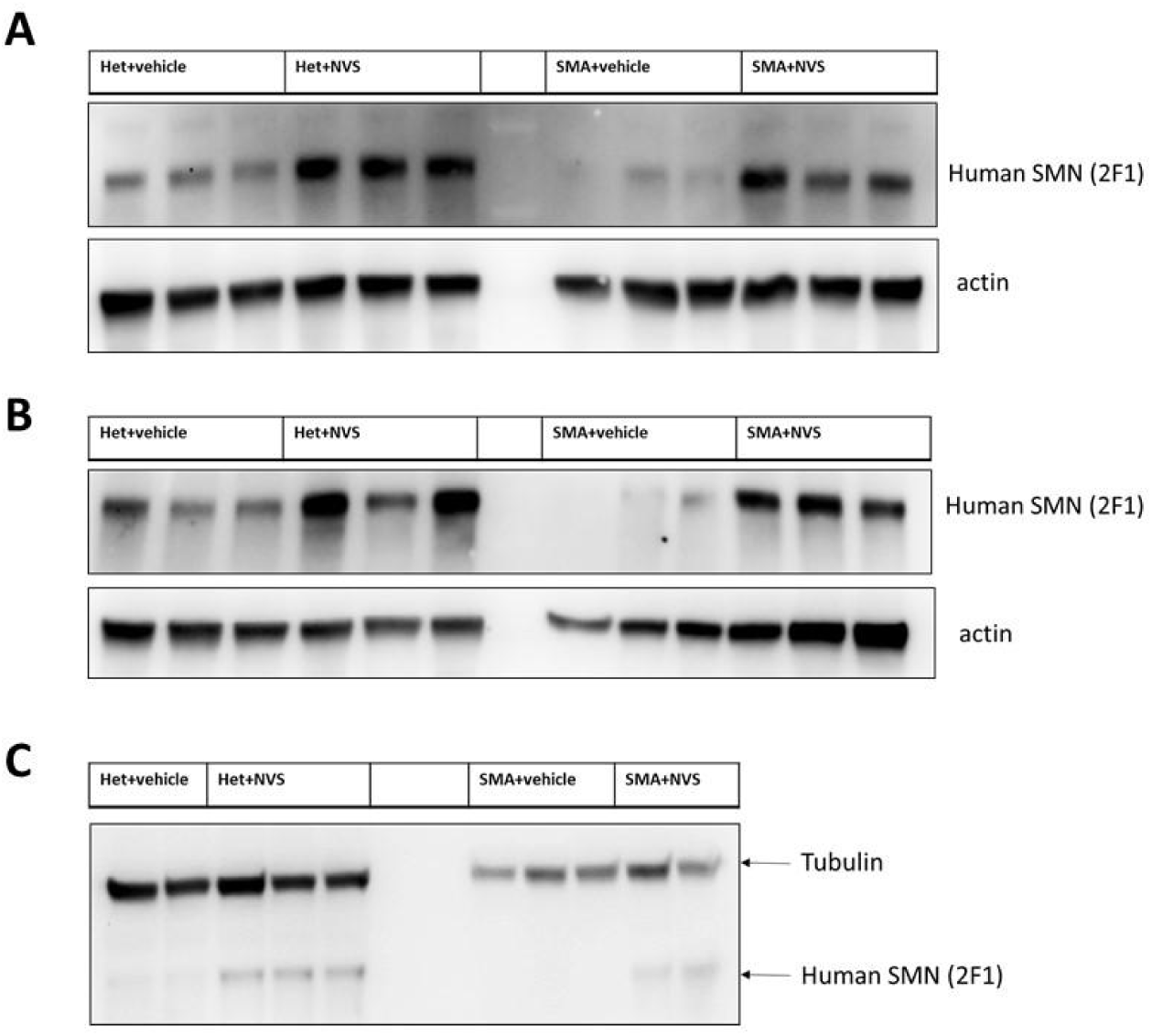
NVS-SM2 increases SMN protein in severe 5058 SMA and control Het mice. Mice were injected subcutaneously with 1 mg/kg NVS-SM2 or vehicle starting PND 2 daily until PND 6. Mice were sacrificed and tissues harvested on PND 7. Human SMN protein levels were analyzed in brain (A), spinal cord (B) and muscle (C) tissue via immunoblotting. SMN protein was normalized to actin and tubulin. Each lane represents tissue from an individual mouse. SMA represents the severe 5058 SMA mice, which express the human SMN2 transgene, and Het mice are their littermates, expressing both human and mouse SMN.

As SMN protein levels increased in the central nervous system and the periphery, we evaluated the biological impact of NVS-SM2 on the temporal development and progression of the SMA phenotype. The severe SMA mice were injected s.c. with NVS-SM2 at 0.1 and 1 mg/kg daily from PND 2-15, followed by every other day until PND 30. All mice were alive at PND 30, demonstrating a successful intervention as severe SMA mice typically succumb by PND 11. Hence, we stopped compound delivery and continued monitoring the mice for lifespan and any phenotypic changes. We observed a median survival of PND 57 and PND 94 after a 30-day treatment with 0.1 and 1 mg/kg, respectively (Figure 2A). The surviving mice were mobile, but progressive distal limb, tail, and ear necrosis necessitated euthanasia (Supplemental video 1). Kaplan-Meier survival curves with the Mantel-Cox log rank test demonstrated that both treatment dosages significantly differed from the vehicle treatment (VH vs. 0.1 mg/kg: p= 0.0036; VH vs. 1 mg/kg: p=0.0014). Heterozygous control mice and NVS-SM2-treated severe SMA mice weighed ∼20 g by PND 30, and weight began to decline at PND 63 in the 1 mg/kg-treated severe SMA group (Figure 2B). Ear necrosis emerged at PND 73. To statistically compare the phenotype between groups, we chose the time point at which the treatment group gained its maximum average weight (MAW). For the 1 mg/kg treatment group, the MAW was at PND 63 with no significant difference in weight between severe SMA mice and Het controls. As a phenotypic marker, we also measured tail length, which was slightly shorter in untreated SMA mice compared with Het mice, and correlated with body weight (Figure 2C, Supplemental Figure 3A). Tail length in the 30-day 1 mg/kg group was slightly shorter than in the Het group, but the difference was not significant (PND 63: 1 mg/kg vs. Het vs.: 7.33 ± 0.17 vs. 8.13 ± 0.24 cm; p=0.054, Figure 2D). The MAW for the 0.1 mg/kg group was reached at PND 32, and low-dose-treated mice were significantly lighter than age-matched Het controls (PND 32: 0.1 mg/kg vs. Het: 15.23 ± 2.47 g vs. 20.1 ± 0.79; p= 0.0296).

**Figure 2.**
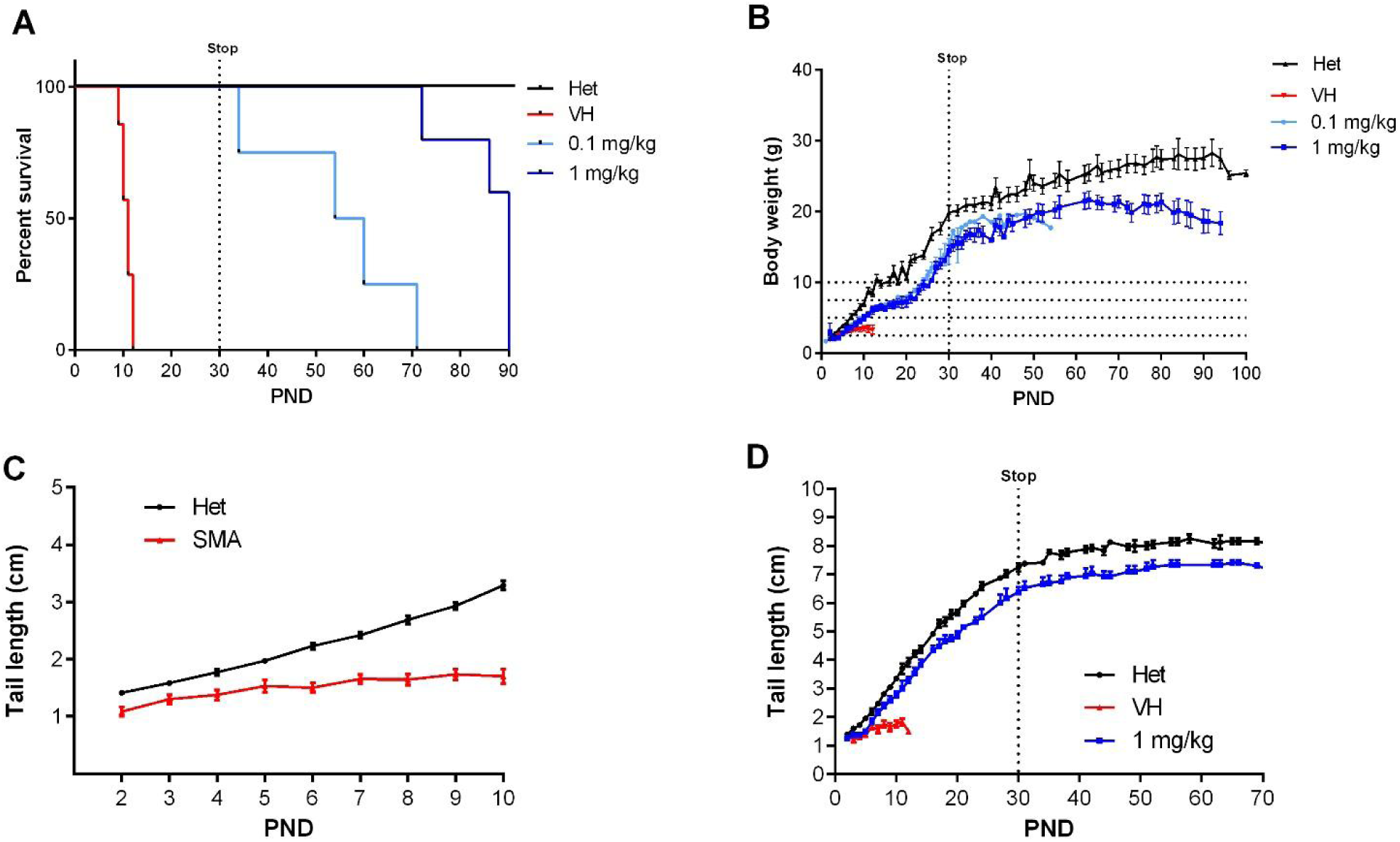
Effect of different doses of NVS-SM2 on severe 5058 SMA mice. Kaplan-Meier survival curves of severe 5058 SMA mice subcutaneously treated with vehicle or 0.1 (n=4) and 1 mg/kg (n=5) NVS-SM2 daily until PND 15 and then every other day until PND 30 (A). Mice were monitored for body weights (B) and tail length (C,D).Data expressed as S.E.M.

This optimized low-dose 30-day treatment regimen represents a tractable mild SMA mouse model that resembles the phenotypic delay in human Type II/III SMA patients. Both doses of subcutaneous NVS-SM2 greatly improved phenotypic outcomes compared with oral administration of NVS-SM1 in SMNΔ7 mice (Palacino *et al*., 2015). Daily oral administration of NVS-SM1 at 3 mg/kg was reported to rescue 60% of SMA mice at PND 30. However, we predict that treatment with 1mg/kg NVS-SM2 past PND 30 will yield the same body weight and tail length between severe SMA and Het mice. Additionally, continued treatment at 0.1 mg/kg may suffice for full rescue.

Based on this unexpected long-term rescue with very low doses of NVS-SM2, we investigated whether a shorter treatment duration would result in a moderate SMA model in severe SMA mice. Neonatal mice were injected once daily with 1 mg/kg s.c. NVS-SM2 on three consecutive days (PND 2-4). Surprisingly, the median survival increased to 30 days, and body weight averaged 10 g, approximately 70% of age-matched Het control weight (Figure 3A, B). The MAW at PND 25 and was significantly different between Het vs. 1 mg/kg (Het vs. 1 mg/kg: 10.5 ± 0.67 vs. 15.4 ± 0.47 g; p= 0.0002). We then decreased the concentration of NVS-SM2 to 0.1 and 0.5 mg/kg and repeated the treatment regimen. These mice showed a dose-dependent decline in median survival to PND 18.5, which is 2.5 days before weaning, and PND 26, respectively. The survival curves of all treatment groups were significantly different from untreated control severe SMA mice (VH vs. 0.1 mg/kg: p=0.0004; VH vs. 0.5 mg/kg: p= 0.0004; VH vs. 1 mg/kg: p=0.0004, Figure 3A). Mice injected with the lowest dose exhibited the lowest gain in body weight and reached their MAW at PND 14 (PND 14: Het vs. 0.1 mg/kg: 9.9±0.3 vs. 6.7±0.4 g; p< 0.0001; Figure 3B), whereas the 0.5 mg/kg-treated mice reached their MAW at PND 22 (PND 22: Het vs. 0.5 m/kg: 12.3±0.3 vs. 8.9±0.3; p< 0.0001; Figure 3B). To test short-term treatment with an even further reduced drug amount, we injected 0.1 mg/kg s.c. for two days - PND 2 and PND 3. This introduced greater variability, with a median survival of 13 days and one mouse surviving until PND 24, rendering this treatment schedule unsuitable (Figure 4A). Body weights were only marginally improved (Figure 4B).

**Figure 3.**
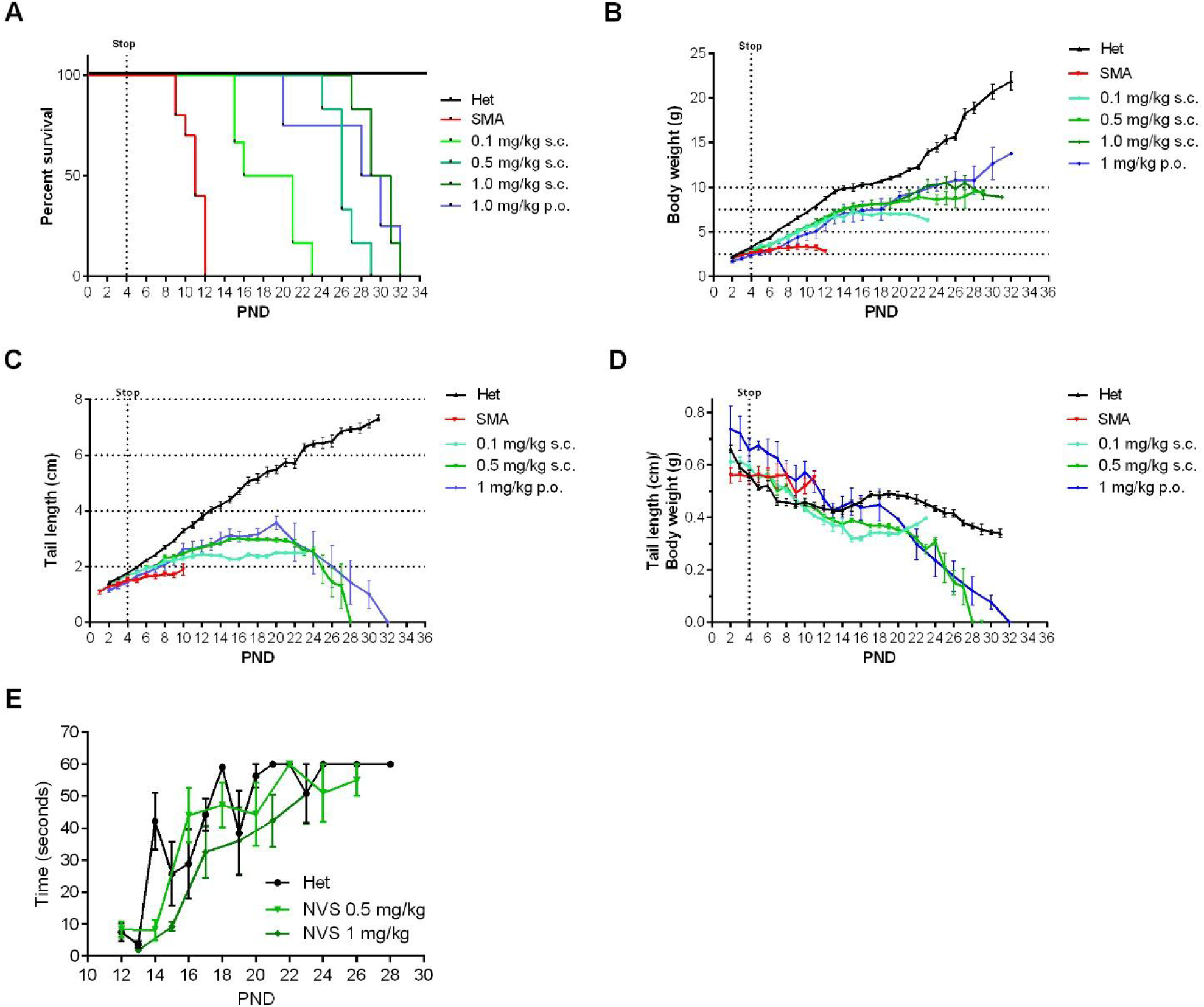
Effect of 3-day treatments with different dosages of NVS-SM2 on severe 5058 SMA mice. Severe 5058 SMA mice were treated on PND 2, PND 3 and PND 4 subcutaneously (s.c) or orally (p.o.) with the indicated doses of NVS-SM2. A) Kaplan-Meier survival curve of severe 5058 SMA mice subcutaneously treated with 0.1 (n=6), 0.5 (n=6), and 1 mg/kg (n=6) NVS-SM2 or orally with 1 mg/kg NVS-SM2 (n=4) and untreated SMA mice (n=10). Mice were monitored for body weights (B) and tail length (C) to determine the tail length: body weight ratio (D). Pen test time of 0.5 mg/kg NVS-SM2 treated mice in comparison to Het mice (E). Data expressed as S.E.M..

**Figure 4.**
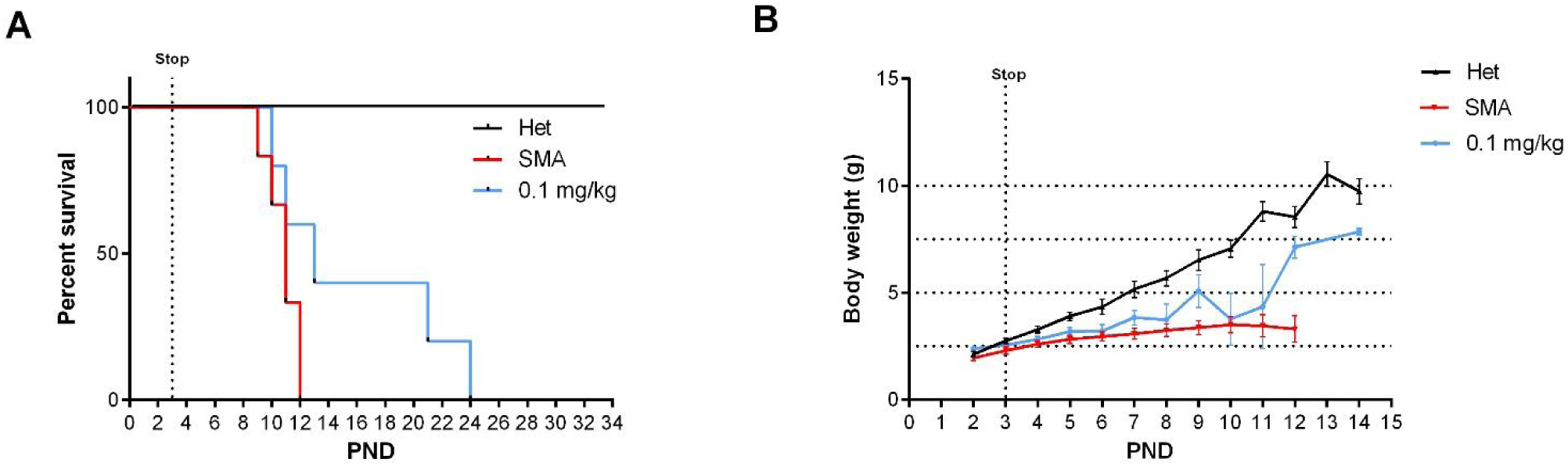
Impact of NVS-SM2 treatment on severe 5058 SMA mice survival after 2-day treatments. Severe SMA mice were treated on PND 2 and PND 3 with the indicated doses of NVS-SM2. A) Kaplan-Meier survival curve of severe 5058 SMA mice subcutaneously treated with 0.1 mg/kg NVS-SM2 (n=5) on PND 2 and PND 3 in comparison to untreated SMA mice (n=6). C) Mice were monitored for body weights. Data expressed as S.E.M.

To investigate whether NVS-SM2 is effective when administered orally, we treated mice with 1 mg/kg for three days (PND 2-4). These mice had a median life span of 29 days (VH vs. 1 mg/kg p.o.: p=0.0024, Figure 3A). The MAW was reached at PND 18 with an average weight of 7.5±1.2 g. This included a runt with a weight of 1.1 g on PND 2 that grew to 4 g at PND 18, while the remaining treated severe 5058 SMA mice had an average weight of 1.9 g on PND 2 and reached 8.7±0.4 g on PND 18 (Figure 3B). Despite the low birth weight and the small volume (2.2 µl) of NVS-SM2 administered, the successful response to the oral drug delivery was remarkable. Tail length in the 0.5 mg/kg group peaked at PND 16 at 3 cm, and tails were much thinner. Tail necrosis followed tail thickening and was overt at PND 23. Tail length of mice treated with the lowest dose peaked at PND 14 at 2.4 cm. These animals were found dead before appearance of tail necrosis (Figure 3C, D), though their tails were notably thinner. This coincides with the observation that the tails of 5058 SMA mice with two copies of *SMN2* become necrotic after weaning (PND 21). Although the ratio of tail length to body weight was not significantly different in severe SMA vs. Het mice (Supplemental Figure 3A), there was a significant difference in the three-day treatment groups (0.5 mg/kg s.c. and 1 mg/kg PO) compared with Het mice (Figure 3D), revealing tail length as a useful and early phenotypic marker of rescue. Mice were analyzed for gross motor function using the beam/pen test starting at PND 12. On average, treated severe 5058 SMA mice performed as well as Het control mice (Figure 3E).

Since SMA research is commonly conducted using the SMNΔ7 transgenic mouse strain, we also investigated the outcome of the three-day treatment regimen in these mice. NVS-SM2 was injected once daily s.c. at 1 mg/kg on PND 2-4, resulting in a median survival of 29.5 days compared with 14.5 days in vehicle-treated mice (p= 0.004, Figure 5A). Body weights were improved and peaked at PND 16 (Figure 5B). These results are comparable to those observed in NVS-SM2-treated severe SMA (5058) mice, demonstrating this approach to be suitable across SMA mouse models. The impressive potency of NVS-SM2 appears to surpass NVS-SM1, which rescued ∼60% of SMNΔ7 mice after daily oral administration at 3 mg/kg (Palacino *et al*., 2015).

**Figure 5.**
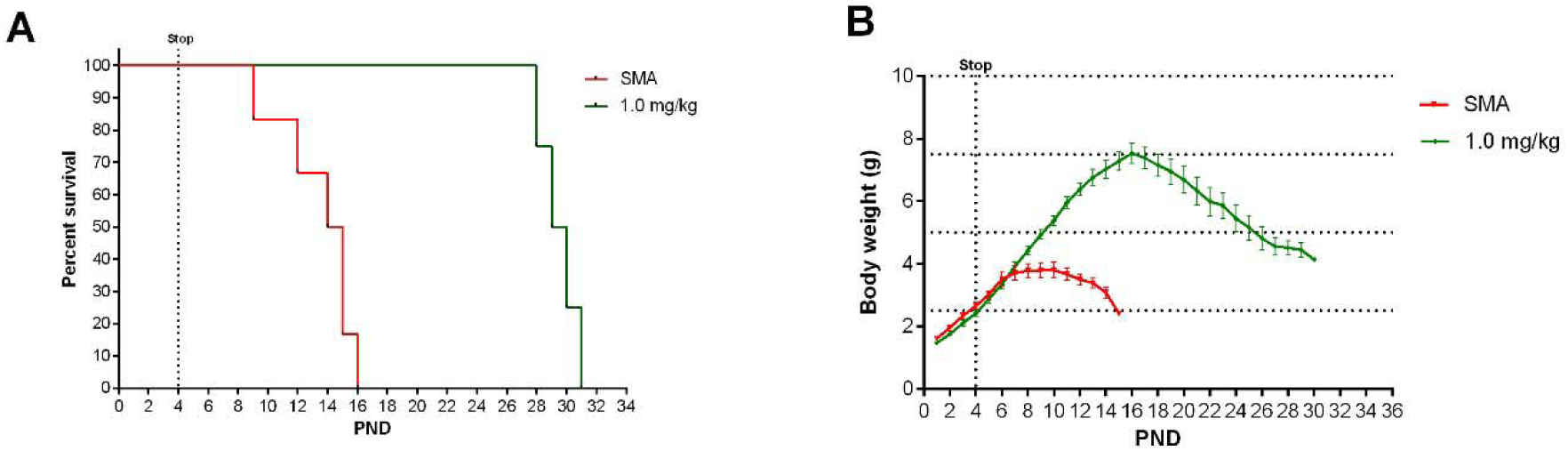
Effect of 3-day treatments with NVS-SM2 on SMNΔ7 mice. SMNΔ7 mice were treated on PND 2, PND 3 and PND 4 subcutaneously with 1 mg/kg NVS-SM2. A) Kaplan-Meier survival curve (A) and body weights (B) of untreated (n=10) and NVS-SM2 (n=4) treated SMNΔ7 mice. Data expressed as S.E.M..

Due to the *in vivo* potency of NVS-SM2, we investigated next the suitability of NVS-SM2 to develop an animal model allowing for the initiation of therapeutic intervention during the symptomatic stage. The majority of the post-symptomatic rescue experiments in severe SMA mice have been explored with antisense oligonucleotides (ASO) and disease-modifying gene therapies (Foust *et al*, 2010; Gogliotti *et al*, 2012; Le *et al*, 2011; Robbins *et al*, 2014). The critical therapeutic window in SMNΔ7 mice is PND 5-6(Foust *et al*., 2010; Gogliotti *et al*., 2012; Le *et al*., 2011; Robbins *et al*., 2014). SMNΔ7 mice, on average, survive four days longer than severe 5058 SMA mice. SMNΔ7 mice injected intracerebroventricular (ICV) with scAAV9-SMN on PND 8 had a median survival of 18 days compared with 14 days in non-injected mice. Mice injected a day earlier showed a greater response with a median survival of 28 days, and a few mice survived for 70 days (Robbins *et al*., 2014). Intravenous injection of scAAV9-SMN on PND 5 modestly increased survival (30 days). Injection at PND 10 did not rescue early lethality. Inducible transgenic rescue experiments also point to PND 6 as the latest time point to rescue SMNΔ7 mice (Le *et al*., 2011; Lutz *et al*, 2011). However, a study in a pharmacologically induced mild SMNΔ7 mice model reported that mice treated with suboptimal dosing of SMN-C3 changed to optimal dosing at a late stage (PND 32) had increased body weight and improved phenotypic markers compared with mice that continued to receive the suboptimal dose in mice (Feng *et al*., 2016).

Few post-symptomatic rescue experiments have been conducted in severe 5058 SMA mice. One study showed treatment (s.c.) with ASOs early in disease resulted in substantial rescue, whereas treatment at PND 5-7 increased survival only to PND 16 (Hua *et al*, 2011). The maturation of the neonatal blood-brain-barrier may restrict ASO access. We performed a trial experiment with NVS-SM2 s.c. at 1 mg/kg starting at PND 6, a time point considered symptomatic in 5058 SMA mice (Groen *et al*, 2018). Mice received daily drug injections until PND 15, then every other day until PND 30, at which point treatment was stopped. Mice had a median survival of 70 days, slightly lower than the median survival of 94 days achieved with PND 2 treatment (p= 0.0063, Figure 6A). Therefore, we investigated if long term survival could be achieved at late symptomatic stages of the disease. We chose PND 8 as the last feasible time point since severe 5058 SMA mice have a median life span of 10.5 days, with some living only until PND 9 and rendering PND 9 as an impractical starting point. Groen EJN et al. showed that at PND 8 SMN protein is decreased systemically by 2-3 fold compared to symptomatic (PND 5) and control litter mates (Groen *et al*., 2018). Mice were injected with 1 mg/kg, s.c. once per day beginning at PND 8. This protocol generated two distinct groups: short-term and long-term survivors. The mice that did not respond succumbed with a median life span of 12 days; survivors continued to thrive (Supplemental video 2), and injections were continued four times per week. The experiment was stopped at PND 110 (Figure 6B). The body weight of the surviving NVS-SM2-treated mice averaged approximately 90% of age-matched Het controls (Figure 6C). The tail was significantly shorter in all survivors (Figure 6D). Body weight at treatment start (PND 8) did not influence the outcome (Figure 7B). Small spleen size is a hallmark of the SMA phenotype in mice (Khairallah *et al*., 2017) and spleen weights were lighter than control spleens, though the difference was not significant (SMA vs. Het: 89.2 ± 8.1 mg vs. 104 ± 3.8 mg) and no difference was observed when spleen weight was normalized to body weight (Figure 7A). SMN protein levels in the brain and spinal cord in long-term survivors were similar to that of control Het mice (Figure 8 A-D). Our study, together with Risdiplam™ data, indicates the potential benefit of splicing-modifier drugs later in disease in both severe and mild SMA. In addition, we show that there is a treatment window beyond PND 7 in the severe 5058 SMA mice. Additional regimens with initial three-day drug exposure and later stage pulses can be considered to evaluate disease progression and the rescue capabilities of NVS-SM2, using weight gain, tail length, and the emergence of digital necrosis as indicators of disease progression.

**Figure 6.**
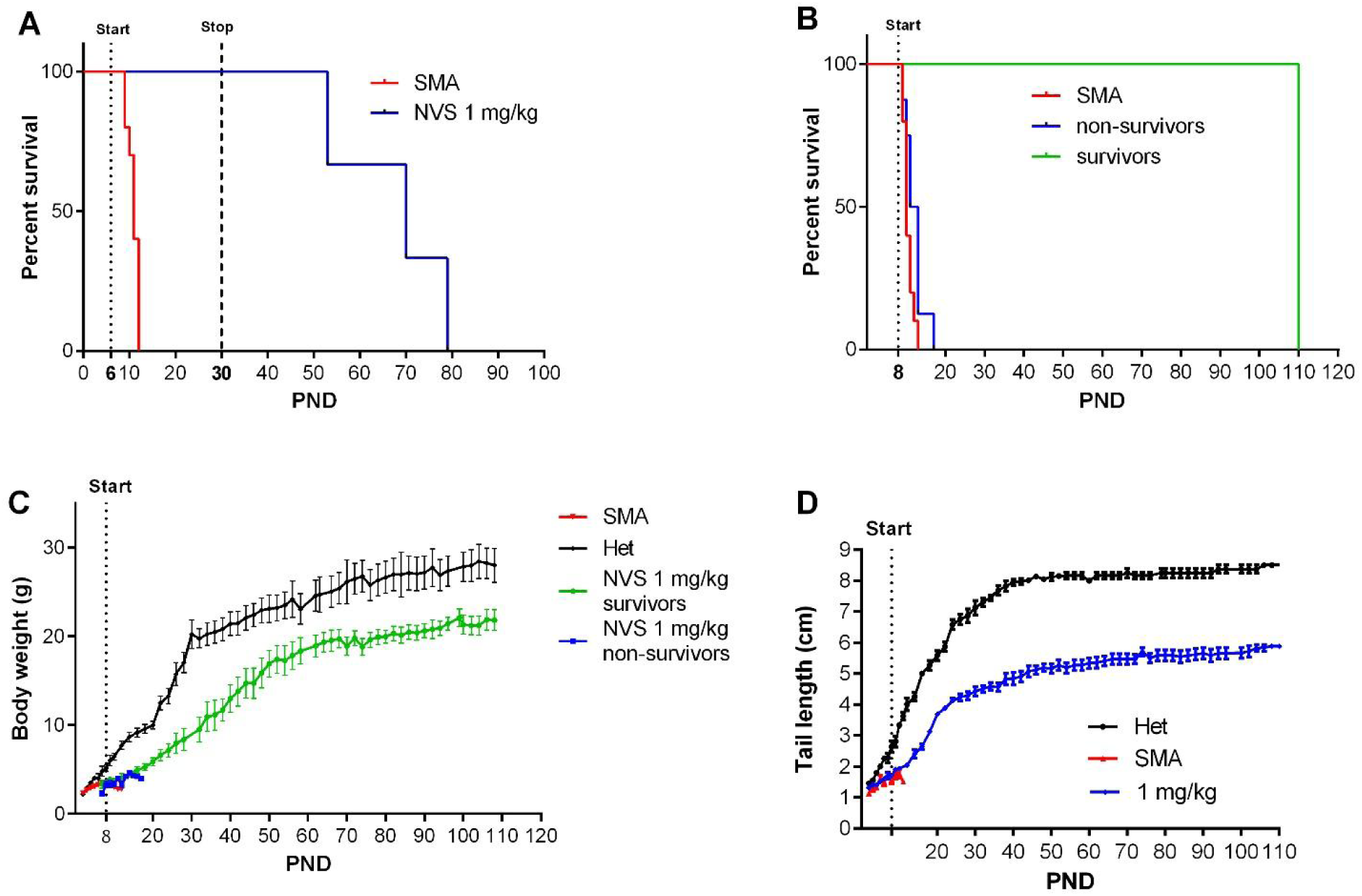
Effect of late treatment of severe SMA (5058) mice with NVS-SM2. Animals were treated starting on PND 6 (A, n=3) or PND 8 (B, n=14) subcutaneously with 1 mg/kg NVS-SM2. A, B) Kaplan-Meier survival curves. PND 8 treated SMA mice were further separated into non-survivors (n=8) and survivors (n=6) groups. PND 8 treated severe 5058 SMA mice were monitored for body weights (C) and tail lengths (D). Data expressed as S.E.M..

**Figure 7.**
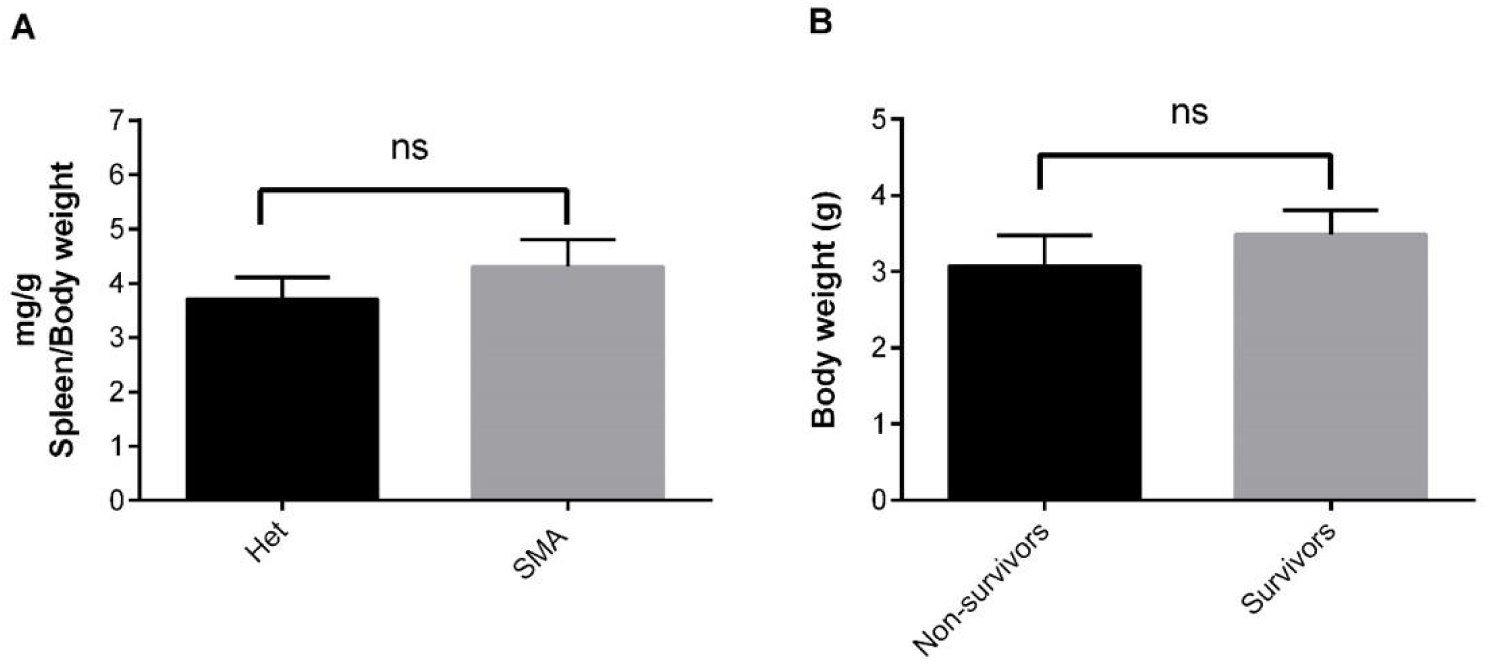
Effect of late treatment of NVS-SM2 in severe 5058 SMA mice. Spleen weights of NVS-SM2 treated severe 5058 SMA mice at PND 110 in comparison to healthy Het control mice are not different (A). Average body weight at PND 8 of the treated NVS-SM2 mice separated into the survivors and non-survivors groups are not different (B). Data expressed as S.E.M. and analyzed using Student’s unpaired t-test, a p-value of 0.05 was taken as significant. ns: non-significant.

**Figure 8.**
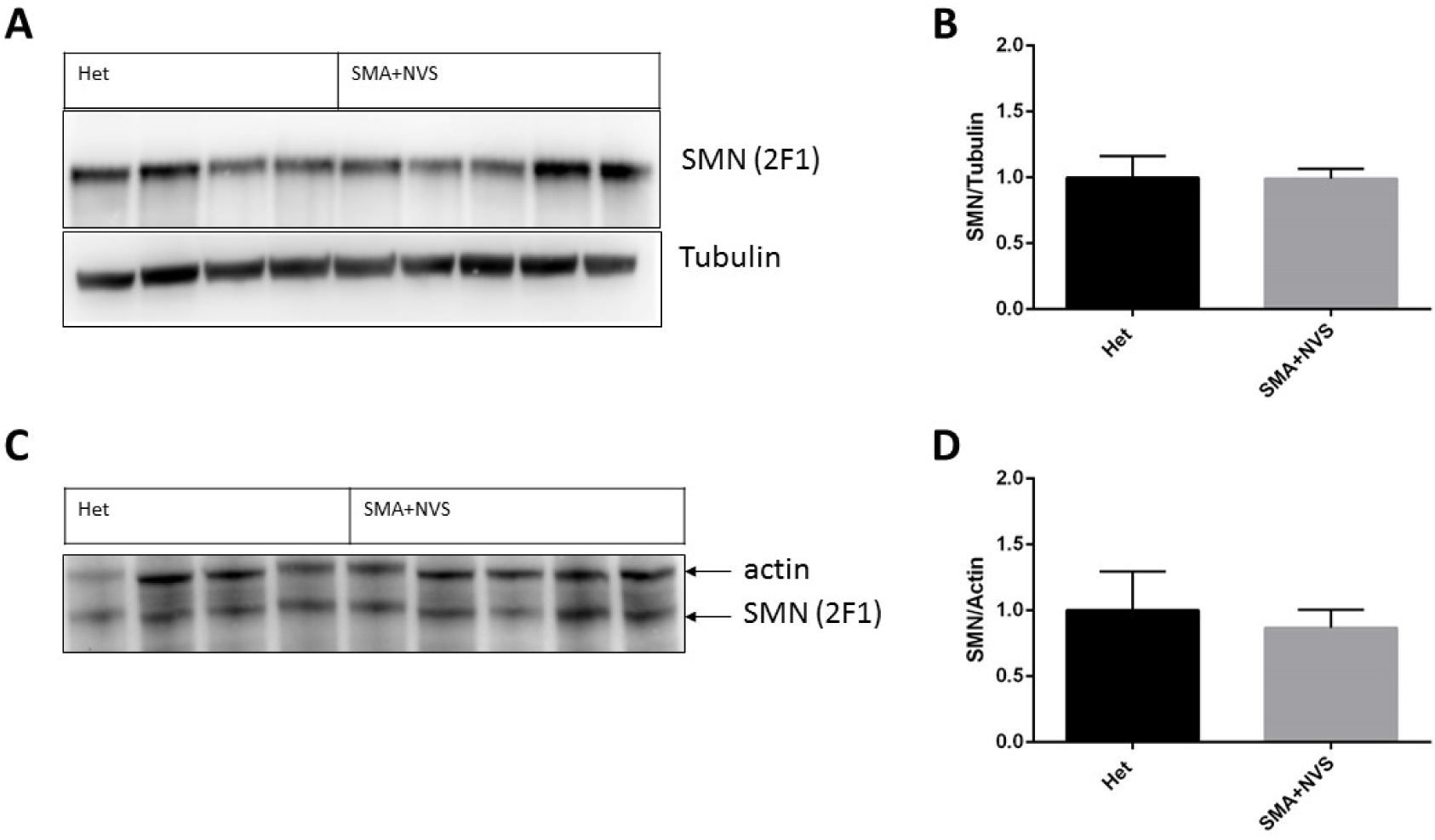
SMN protein levels in NVS-SM2 treated severe 5058 SMA mice at PND 110 in compared to healthy Het control mice. Immunoblot of human SMN and housekeeping proteins at PND 110 in Het (n=4) and NVS-SMN2 PND 8 treated severe SMA 5058 mice (n=5) in brain (A, B) and spinal cord (C, D). SMN protein was normalized to housekeeping proteins (B, D). Each lane represents tissue from an individual mouse. Data expressed as S.E.M. and analyzed using unpaired t-test, a p-value of 0.05 was taken as significant.

In summary, the *SMN2* splicing modifier NVS-SM2 is highly active *in vivo* and could be titrated in dosage, timing, and duration of administration for the development of robust SMA models at varying disease stages. NVS-SM2 increased SMN protein in the brain, spinal cord, and muscle tissues in severe SMA (5058) mice and extended their lifespan and bodyweight. Subcutaneous and oral administration on PND 2-4 is sufficient to extend survival with dose-dependent responses. The limited invasiveness of the three-day treatment model eases the stress on the neonatal mice, providing a significant advantage over daily injections. Drug-drug interactions are eliminated due to the short duration of drug administration and the reported short half-life of NVS-SM2. NVS-SM2 rescued also severe SMA mice at late symptomatic time points. Treatment starting at PND 8 resulted in a group that survived until the end of the experiment at PND 110. This treatment starting point has not been therapeutically beneficial with antisense oligonucleotide or adeno-associated virus-9 SMN gene therapy. We speculate that both systemic exposure and the dramatic increase in SMN protein levels contribute to this outcome. Indeed, SMN protein levels in the brain and spinal cord were comparable to Het control mice (Figure 8 A-D). Future studies are required to assess the status of other phenotypic SMA markers such as NMJ and muscle pathology in these late-stage rescued animals and the mechanism underlying their recovery. Of particular importance is to investigate how long the SMN induction by NVS-SM2 lasts since the drug has a half-life of three hours, yet we observed a lifespan extension of 26 days past the last administration. This effect provides a significant opportunity to investigate the temporal requirements of SMN in the pathogenesis of SMA.

## Material and Methods

### Materials and antibodies

PEG400 and DMSO were purchased from Sigma-Aldrich. PBS, DMEM, Pen/strep was purchased from Gibco. Fetal bovine serum was purchased from Peak serum. The following antibodies were used: anti-SMN (1:2000; 2F1; Cell Signaling Technologies and MANSMA 6, 4H2, Developmental Studies Hybridoma Bank, DSHB), anti-tubulin (DM1α, 1:4000, Sigma-Aldrich), anti-beta actin (AC-74, 1:4000, Sigma-Aldrich). MANSMA6 (4H2) was deposited to the DSHB by Morris, G.E. (DSHB Hybridoma Product MANSMA6 (4H2)).

### NVS-SM2preparation

NVS-SM2 2-(6-(methyl(2,2,6,6-tetramethylpiperidin-4-yl)amino)pyridazin-3-yl)-5-(1H-pyrazol-4-yl)phenol) was prepared following the procedures described in WO2014028459. LCMS and ^1^H NMR of the final NVS-SM2 were consistent with its structure and the published data^17^.

### Cell culture and reporter cell assay

SMN2 reporter cells were grown in DMEM containing 10% FBS and 1x Pen/Strep. The assay was completed as previously described(Rietz *et al*, 2017). In brief, cells were seeded in a 96-well plate at 25,000 cells/well. The following day cells were treated with 3-fold serial dilutions of compounds, incubated for 24 hrs, lysed, and analyzed using the Dual-Glo^®^ Luciferase Assay System (Promega).

### Mice breeding, genotyping and treatments

The animal study protocols were approved by the Institutional Animal Care and Use Committee of Indiana University and conform to the Guide for the Care and Use of Laboratory Animals. Study protocols were also approved by the University of Missouri Animal Care and Use Committee as well as the regulations established by the National Institute of Health’s Guide for the Care and Use of Laboratory Animals. Severe SMA (5058) neonatal mice were breed as previously reported(Gogliotti *et al*., 2010). Animals were maintained on a 12-12 hour light-dark cycle with food and water *ad libitum* and were provided with Bed-r’Nest as standard of care. DNA was isolated from tail snips of approximately 0.1-0.15 cm length using the Qiagen DNAeasy Kit. SMN1^tm1^ and SMN2 genotyping was performed as directed by Jackson

Laboratory using their suggested primers. Treatment started on Postnatal day 2 (PND 2) unless otherwise stated, with PND 0 as the day mice were born. All treatments used the same vehicle (PEG400: PBS (50:50)) to solubilize NVS-SM2. Due to the high drug potency, 1 mg/ml was prepared and serially diluted to obtain final stock solutions of 0.1 mg/ml to 0.01 mg/ml. Mice were treated subcutaneously with a final dose of either 0.1 mg/kg, 0.5 mg/kg, or 1 mg/kg via daily subcutaneous injection or oral administration. Subcutaneous administration was performed at 10 µl/g body weight, and oral administration at 2 µl/g body weight. Oral administration in neonates was performed using a plastic feeding tube (FTP-20-30-50, Instech Laboratories). Feeding tubes were placed in the inner-cheek, and the suspension was applied slowly, stimulating suckling behavior. In brief, the following treatment schedules were used a) 30-day treatments started on PND 2, everyday s.c. until PND 15 followed by every other day until PND 30, b) 3-day treatment groups received treatment on PND 2, PND 3, and PND 4 with the indicated doses c) 2-day treatment groups on PND 2 and PND 3 d) PND 6 late treatment: starting PND 6, daily until PND 15 from then followed by every other day until PND 30, and e) PND 8 late treatment: starting PND 8, daily until PND 15 from then four times a week until PND 110. Mice were euthanized via CO2 exposure and the whole brain, spinal cord, spleen, and muscle (left vastus lateralis) were extracted for western blotting. We used the balance beam test to assess motor balance and coordination in treated SMA mice, as described in SMA_M.2.1.001 published by TREAT-NMD Neuromuscular Network. The beam/pen test was conducted every other day from PND 12 until PND 30. For the severe SMNΔ7 animal model, heterozygous breeder pairs of mice (Smn+/-;SMN2+/+;SmnΔ7+/+), were purchased from the JAX^®^ Laboratory (JAX^®^Stock#005025:FVB.CgGrm7Tg (SMN2)89AhmbSmn1tm1MsdTg(SMN2*delta7)4299Ahmb/J; The Jackson Laboratory, 610 Main Street Bar Harbor, ME 04609 USA). The colony was maintained as heterozygote breeding pairs under specific pathogen free conditions. Experimental mice litters (Smn-/-;SMN2+/+;SMNΔ7+/+ referred as SMNΔ7) were genotyped on the day of birth (PND0) using standard PCR protocol (JAX^®^ Mice Resources) on tail tissue material as previously described. Experimental pups were kept with a minimum of two healthy heterozygous siblings.

### Tissue isolation and Immunoblotting

Extracted whole brain, spinal cord, and vastus lateralis were extracted on PND 7 and lysed in the pre-heated lysis buffer (2% SDS, 150 mM NaCl, 10 mM Tris-HCl, pH 8.0 + 1x Pierce protease inhibitor cocktail (added fresh, A32961) at a tissue:buffer volume ratio of 1:15. Tissues were heated in lysis buffer for 5 min at 95°C. Brain and spinal cord were disrupted and homogenized using a 22g needle, while muscle tissue was first homogenized using an 18g needle followed by a 22g needle. Lysates were then heated for 10 min at 95°C, and cleared by centrifugation at 8000xg for 10 min. Protein concentration was measured using the Pierce BCA kit, and equal amounts of protein were separated on a 4-12% SDS-gel (Genscript SurePAGE™). Proteins were transferred onto a PVDF Immobilon-P (0.45 µm Millipore), and SMN, b-actin, and a-Tubulin proteins were visualized by chemiluminescence after exposure to the anti-SMN antibody (Cell Signaling 2F1) following by an HRP linked secondary antibody. Signal intensities were quantified using ImageJ and Image Studio™ Lite (LI-COR Biosciences). SMN expression is expressed as fold change and normalized to housekeeping protein, b-actin, and a-Tubulin as indicated.

### Data analysis and statistics

Survival was analyzed with Kaplan-Meier survival curves using the log-rank Mantel-Cox test for survival comparisons. (Graph-Pad Prism v6.00; GraphPad Software, Inc.). A p-value of p<0.05 was considered statistically significant. The maximum average weight (MAW) is defined as the maximum average weight at the last time point at which all mice of the treatment group that entered the study were still alive. This time point was used to determine if body weights or tail length differences were statistically different. Groups of two were analyzed using unpaired t-test, and groups and groups of more than two were analyzed using one-way ANOVA with post-hoc analysis (Dunnett’s or Bonferroni as indicated). All data are expressed as standard error of the mean (S.E.M.) unless otherwise stated.

## Acknowledgments

This research was supported by grants NINDS R33NS095139 and CureSMA to EJA, CLL, and KJH. The content is solely the responsibility of the authors. We thank Jacob Astroski for assistance with mouse experiments, and Academia Sinica for providing the 5058 mice.

